# Global alignment and local curvature of microtubules in mouse fibroblasts are robust against perturbations of vimentin and actin

**DOI:** 10.1101/2023.04.19.537509

**Authors:** Anna Blob, David Ventzke, Ulrike Rölleke, Giacomo Nies, Axel Munk, Laura Schaedel, Sarah Köster

## Abstract

The eukaryotic cytoskeleton is an intricate network of three types of mechanically distinct biopolymers – actin filaments, microtubules and intermediate filaments (IFs). These filamentous networks determine essential cellular functions and properties. Among them, microtubules are important for intracellular transport and establishing cell polarity during migration. Despite their intrinsic stiffness, they exhibit characteristic bending and buckling in cells due to non thermal forces acting on them. Interactions between cytoskeletal filaments have been found but are complex and diverse with respect to their effect on the mechanical behavior of the filaments and the architecture of networks. We systematically study how actin and vimentin IFs influence the network structure and local bending of microtubules by analyzing fluorescence microscopy images of mouse fibroblasts on protein micropatterns. Our automated analysis averages over large amounts of data to mitigate the effect of the considerable natural variance in biological cell data. We find that the radial orientation of microtubules in circular cells is robust and is established independently of vimentin and actin networks. Observing the local curvature of microtubules, we find highly similar average bending of microtubules in the entire cell regardless of the cytoskeletal surrounding. Small systematic differences cannot be attributed directly to vimentin and actin densities. Our results suggest that, on average, microtubules in unpolarized mouse fibroblasts are unexpectedly independent of the rest of the cytoskeleton in their global network structure and their local response to forces.

## Introduction

The cytoskeleton of eukaryotes is an intricate network comprising three different biopolymers: microtubules, actin filaments and intermediate filaments (IFs). Each of these cytoskeletal filament types has their own distinct properties and forms unique architectures within the cell.^1^ Together, they are essential for various cellular processes, such as cell division, intracellular transport, migration and maintaining the mechanical integrity of the cell. ^2–5^ Microtubules often show a radial network organization in motile cells with microtubules growing outwards from the centrosome or the Golgi complex.^6,7^ Since microtubules support cell protrusions and form tracks along which motors can transport cargo such as organelles and vesicles, a robust microtubule network organization is important for, e.g., directed cell migration and the cell’s ability to dynamically adapt its polarization.^5,8^ Microtubules are embedded in a crowded and mechanically challenging environment, containing – among other components – actin and vimentin networks. This raises the question of what ensures the reliable development and robustness of a reproducible microtubule organization in the cell. Examples of microtubules being guided by other cytoskeletal filaments have been found: Actin fibers lead microtubules to focal adhesions^9,10^ and vimentin IFs template the microtubule network in migrating RPE1 cells, thus helping to maintain the polarity of the cell. ^11^ It is not completely clear, however, to what extent vimentin and actin networks are required for microtubules to be oriented from the center to the periphery and their overall network organization.

On a more local scale, the intrinsically stiff microtubules with a persistence length on the millimeter scale^12^ show strong bending in cells which is attributed to nonthermal forces acting on them.^13–16^ When describing a single microtubule as a beam that is compressed, it is considered to be influenced by its elastic surrounding.^17,18^ In vitro, without an elastic support, a microtubule displays buckling in the shape of a single arc, but if a beam is embedded in an elastic matrix it buckles with a much shorter wavelength depending on the elastic modulus of the surrounding.^18^ The elastic modulus of the whole cell and of the cytoplasm has been found to depend on actin and vimentin networks, which generally make a cell stiffer.^19–22^ The extent to which this notion of individual, buckling microtubules in an elastic cytoskeletal matrix can be transferred to the entire ensemble of microtubules with their various shapes is not clear. Previous studies with contrasting results present an inhomogeneous and complex picture of the influence of the cytoskeletal surrounding on microtubule shapes. For instance, for individual dynamically buckling microtubules in Cos7 cells an increase of buckling wavelength was found when depleting actin^18^ thus confirming the constrained buckling theory of elastically supported microtubules and in cryotomograms of HeLa and P19 cells, a lower persistence length was found for microtubules that appear to be entangled with actin and vimentin filaments.^23^ In contrast, the authors of a study of microtubule bending in living Xenopus laevis melanophores observed a smaller persistence length when disrupting vimentin IFs.^24^

Here, we present a systematic study of the influence of vimentin IFs and actin filaments on microtubules in mouse embryonic fibroblasts (MEFs) on circular patterns. This cell type is selected for its wide-spread use in a variety of studies and for its biological importance as a typical motile cell of mesenchymal origin.^22,25,26^ Microtubules in cells with and without vimentin and/or actin filaments are analyzed in an automated way allowing us to analyze large amounts of data and to distinguish average behaviors from natural cell-to-cell variability. We find that the alignment and orientation of microtubules in unpolarized mouse fibroblasts are robust against the lack of vimentin and against perturbations of actin. The average local curvature of microtubules shows only small differences for cells or cell areas with and without actin or vimentin hinting to an indirect effect rather than a direct causality between microtubule curvature and vimentin or actin network density. Our study highlights the importance of a bias-free analysis to extract average behavior and shows that the organization and shape of microtubules in unpolarized mouse fibroblasts may be surprisingly independent of their cytoskeletal surrounding.

## Materials and methods

### Micropatterning

Fibronectin micropatterns are prepared on glass coverslips (No. 1.5H, Carl Roth, Karlsruhe, Germany) or glass bottom dishes (Ibidi, Gräfelfing, Germany, #81218-200) using the photopatterning system PRIMO (Alvéole, Paris, France) mounted on an inverted fluorescence microscope (IX83, Olympus, Tokyo, Japan) equipped with a 20x/NA 0.45 objective (LUCPLFLN 20x, Olympus) and following instructions by Alvéole. Briefly, to passivate the surface, the glass bottom dishes and glass coverslips are treated with air plasma (0.5 mbar at 50 W for 200s, ZEPTO, Diener Electronics GmbH, Ebhausen Germany) and are incubated in two separate steps with 0.2 mg/mL poly-L-lysine (PLL, Sigma, #5899 or #P8920) in ultrapure water for roughly 30 min and with PEG-SVA (mPEG-Succinimidyl Valerate, Laysan Bio, Inc., Arab, Alabama, USA, #MPEG-SVA-5000) in a concentration of roughly 70 mg/mL in 100 mM HEPES buffer, pH 8.1-8.3 for about 60 min. After each incubation, the samples are washed with ultrapure water and dried. For patterning, PLPP gel (Alvéole) is mixed with 70% ethanol and is spread and dried on the coverslip or glass bottom dish. Upon UV-illumination via the PRIMO module, this photo-initiator degrades the PLL-PEG passivation of the glass. The areas to be patterned (in our case circles with a surface area of roughly 1800 *µ*m^2^) are illuminated with a UV dose of 30 mJ/mm^2^. After washing and rehydrating with Dulbecco’s phosphate-buffered saline (PBS), the samples are incubated with fibronectin (from bovine plasma, Sigma-Aldrich, #F1141) at a final concentration of 25-38 *µ*g/mL in PBS for 5 to 15 min. Unbound fibronectin is washed off with PBS.

### Cell culture and drug treatment

Wildtype (WT) MEFs with vimentin and vimentin knockout (VimKO) MEFs without vimentin^27^ are kindly provided by John Eriksson (Åbo Akademi University, Turku, Finland).^25,28^ All cell lines are cultured in high glucose Dulbecco’s Modified Eagle’s Medium (Sigma-Aldrich, #D6429) supplemented with 10% of a serum substitute for fetal bovine serum (Fetal+, Serana, Pessin, Germany, #S-FBSP-EU-015), 1% GlutaMAX (Gibco, Thermo Fisher Scientific) and 1% penicillin-streptomycin (10,000 U/ml penicillin, 10 mg/ml streptomycin, PAN-Biotech, Aidenbach, Germany) at 37^°^C in a humidified incubator with 5% CO_2_. After trypsinization, cells are seeded with a density of 5 000 to 10 000 cells/cm^2^ on fibronectin patterns. After 1 to 2 h, the samples are washed with medium. In total, the cells grow on the patterns for 4 - 5.5h, before they are exposed to a drug or fixed. To freshly polymerize microtubules while actin filaments are disrupted, latrunculin A (Sigma-Aldrich, #428021) at a final concentration of 0.7 *µ*M and nocodazole at a final concentration of 2 *µ*M in cell medium are used. The concentration of DMSO as solvent of the pharmacological agents does not exceed 0.05% in total. After simultaneous incubation with latrunculin A and nocodazole for 20 min, the cells are washed four times with medium which still contains latrunculin A and are further incubated with latrunculin A for 30 min more to let the microtubule network repolymerize.

### Fixation and immunofluorescence staining

Cells on micropatterns on glass are chemically fixed and permeabilized by 4% formaldehyde (methanol-free, Thermo Scientific, #28906), 0.05% glutaraldehyde (Polysciences, Warrington, PA, USA, #01201 or Carl Roth, #4157) and 0.1% Triton X 100 in cytoskeleton buffer (9.3 mM MES, 128.34 mM KCl, 2.79 mM MgCl, 1.86 mM EGTA, adjusted to pH 6.1 with NaOH or KOH) with 9.3% sucrose for 10 min. The samples are rinsed with PBS and subsequently incubated with NaBH_4_ in PBS (∼ 30 mM) for 10 min to reduce background fluorescence. After washing with PBS, the samples are incubated with a blocking solution consisting of 3% bovine serum albumin (BSA; Capricorn Scientific, Ebsdorfergrund, Germany, #BSA-1U or Sigma Aldrich #A7906) and 0.1% Tween 20 (Carl Roth, #9127.1) in PBS for 45 to 60 min at room temperature. Subsequently, the cells are incubated with primary antibodies against vimentin and tubulin diluted in blocking solution at room temperature for about 60 min. After rinsing with PBS containing 0.1% Tween 20, the cells are incubated with secondary antibodies and fluorescently labeled phalloidin in blocking solution at room temperature for about 60 min. Subsequently, the samples are washed again with PBS containing 0.1% Tween 20. Glass coverslip samples are briefly rinsed with ultra pure water and mounted on glass microscope slides with ProLong Glass Antifade Mountant (Invitrogen, Thermo Fisher Scientific, #P36982) whereas for glass bottom dish samples, a clean glass coverslip is mounted on top of the glass well with ProLong Glass Antifade Mountant. As primary antibodies we use mouse monoclonal anti-*α*-tubulin (dilution 1:250, Invitrogen, #62204) and rabbit monoclonal anti-vimentin (D21H3) (dilution 1:200, Cell Signaling Technology, Danvers, MA, USA, #5741). As secondary antibodies we use goat anti-mouse Alexa Fluor 488 (dilution 1:500, Invitrogen, #A-11029) and goat anti-rabbit Alexa Fluor 647 (dilution 1:400, Abcam, Cambridge, UK, #ab150083). For actin staining, we use Alexa Fluor 405 phalloidin (dilution 1:200, Invitrogen, #A30104).

### Microscopy

Fluorescence images are recorded on an inverted laser scanning confocal microscope (Fluoview FV3000 on an IX83 microscope body, Olympus) equipped with a 60x/NA 1.4 oil objective (PLAPON-SC, Olympus) and the following laser lines: 405 nm, 488 nm and 640 nm. The 2D images of each cell are recorded at the lowest z-position within the cell where microtubules are in focus.

### Preprocessing of images

Only images of cells that are fully spread on patterns are considered. Additionally, we choose cells, where the microtubule density is low enough for further processing. From the cells with latrunculin A and nocodazole treatment, only cells without remaining intact actin network and without major detachment from the substrate are selected. All image processing is performed with Python code. Initially, all cell images are aligned such that the circular cell is centered in the image, as described in more detail in the Supplementary information. For presenting the images in figures, they are contrast enhanced. For each category, a binary cell mask is created by segmenting the microtubule image into a cell interior and exterior. This mask is applied in the microtubule alignment analysis to only analyze the alignment of the relevant signal in the cell and not from the signal noise outside of the cell. The mask is also used for dividing each cell individually into an inner center region and an outer rim region. The separation line between both regions is created such that it marks half of the maximum distance to the cell edge.

### Microtubule alignment and orientation analysis

The microtubule images are analyzed with respect to the local degree of alignment of microtubules and their main orientation with Python code based on the AFT tool (Alignment by Fourier Transform).^29^ For each individual image, we subtract the background and normalize by the 95% quantile of the image. To avoid an influence of microscopy artifacts favoring a certain direction, we rotate half of the images by 90^°^ before they are processed. In brief, we calculate a 2D fast Fourier transform (FFT) of overlapping subwindows of an image and perform a principal component analysis (PCA) on the norm of the FFT. With the resulting eigenvalues *λ*_1_ and *λ*_2_ of the covariance matrix of the image, the eccentricity 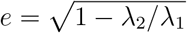 is calculated as a measure for the degree of microtubule alignment with 0 indicating no alignment and 1 indicating maximum alignment. The orientation of the FFT norm and thus of the microtubules is determined as an angle between -90^°^ and 90^°^ with 0^°^ corresponding to a horizontal orientation in the image. Here, this analysis is applied with overlapping windows of size 71×71 pixels that slide along the image with steps of one pixel. For each window, the FFT norms of all cells are averaged before calculating the eccentricity and orientation. The full workflow is depicted in Fig. S3 in the Supplementary information.

### Microtubule filament extraction and curvature analysis

The coordinates of microtubules are extracted with the software SOAX, version 3.7.0.^30,31^ For the processing of the background-corrected microtubule images, we use the default parameters of SOAX and only adjust “ridge-threshold” to 0.03 and “init-z” to false. We choose not to trace filaments across nodes and only consider segments between nodes. For the curvature analysis, only segments longer than 20 pixels (=1.56 *µ*m) are taken into account and their outer three points on each end are omitted to reduce the effect of extraction artifacts in dense node areas. The local curvature of the microtubules is determined by fitting an osculating circle to each filament point taking into account its neighboring points. In total *s* = 13 points contribute with weights that are given by a Gaussian kernel truncated at two standard deviations. Therefore, for the fit to a point at position *i*_0_ the 6 neighboring points to each side at position *i* are given a weight

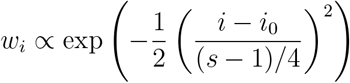

which is then divided by the sum of all weights to rescale the Gaussian kernel to 1. As a consequence, points contribute the less to the fit the further away they are from the point in question. The fit is performed with the least-square method. The inverse of the circle radius corresponds to the curvature.

As an error bar for the mean curvature in the regional analysis, a 95% confidence is constructed as explained in more detail in the Supplementary information.

### Statistical testing

For the statistical comparison of multiple cell categories, a Kruskal-Wallis test is performed. If the result shows a statistically significant difference at a significance level of 5%, a posthoc Dunn’s test with ‘Holm’ level adjustment and a significance level of 5% is performed for pairwise comparisons. Note, that the Holm level adjustment guarantees the overall level of 5% of any wrong rejection in the pairwise comparison.

## Results

To investigate if and how vimentin IFs and actin filaments influence microtubules in cells, we analyze the alignment as well as the local curvature of microtubules in MEFs. We compare cells derived from wildtype (WT) mice and from vimentin-knockout (VimKO) mice^25,27^ to study the effect of vimentin on microtubules. To study the effect of actin, we analyze cells in which the microtubules have freshly polymerized while actin filaments are disrupted. To this end, cells are simultaneously treated with latrunculin A and nocodazole to pharmacologically disrupt both the actin and microtubule network. When nocodazole is subsequently washed out while keeping latrunculin A in the cell medium, the microtubules grow while the actin filaments are still disturbed. In total, we analyze four different categories: WT and VimKO cells each with and without drug treatment. To render individual cells and their cytoskeletal networks comparable, we employ single-cell patterning and thus force the cells to take on a defined circular shape on fibronectin patterns on glass. After chemical fixation and fluorescent staining of the cytoskeleton, confocal fluorescence microscopy images are recorded of the microtubules, the vimentin IFs and the actin filaments. For the four categories WT, VimKO, WT LatA+Noc and VimKO LatA+Noc we record and analyze 82, 65, 61 and 57 cells, respectively. Example images of the three cytoskeletal networks in each of the four cell categories are presented in Fig. 1. As known from previous studies,^32^ microtubules span the entire cell as a network of mostly individual filaments. Actin filaments form dense meshes, bundles and fibers and are primarily located towards the periphery of the cell with occasional prominent stress fibers crossing the entire cell. Vimentin filaments are, just like actin filaments, hardly visible as individual filaments. They tend to bundle and have a rather blurry appearance which we attribute to their lower persistence length. They are typically concentrated around the nucleus and extend from there to the periphery. As expected, we observe a pronounced cell-to-cell variability of each network type. The presented images show individual examples.

**Figure 1.**
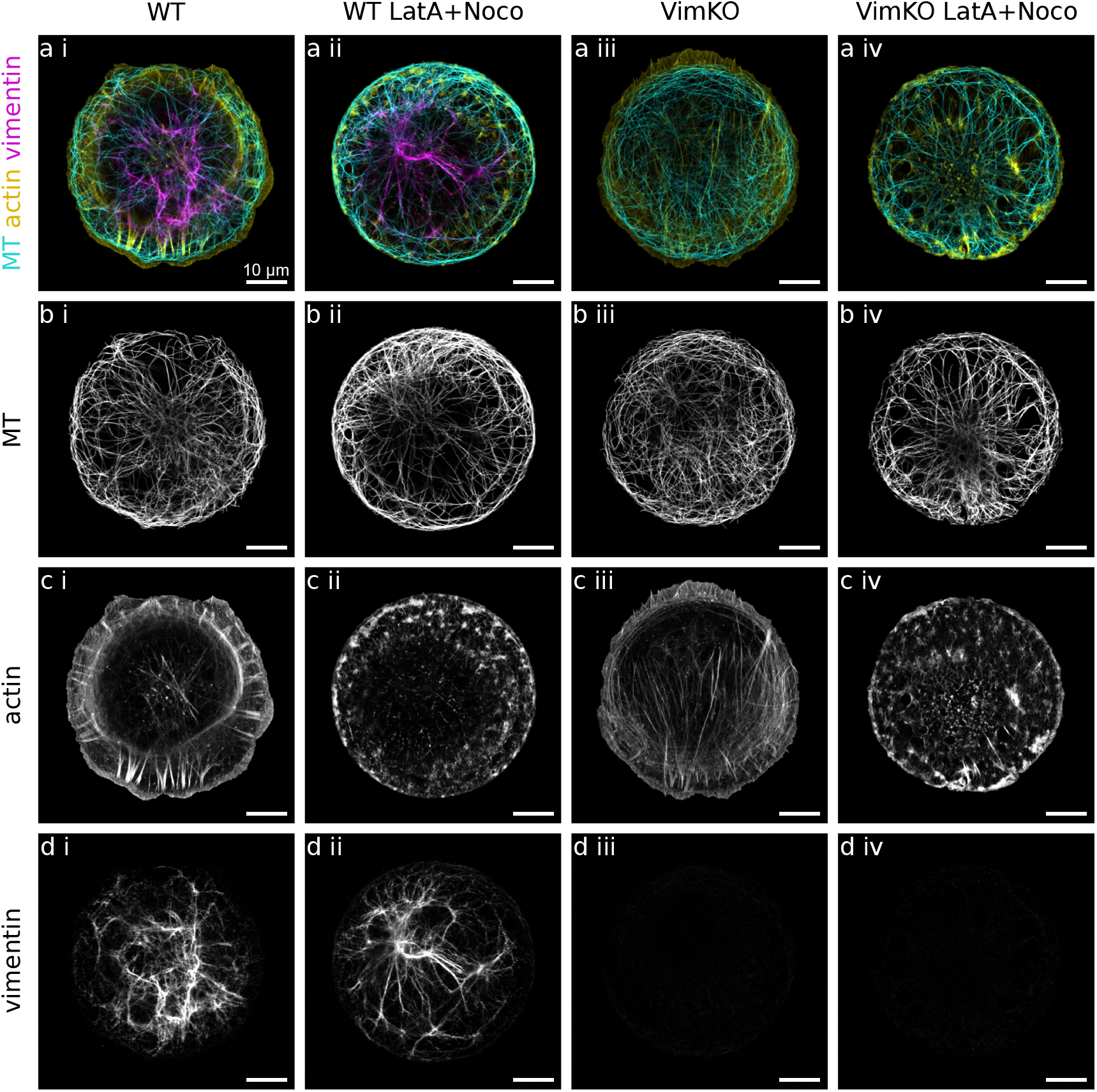
Cytoskeletal networks in cells. Example confocal microscopy images of fixed and fluorescently stained single mouse fibroblasts on circular fibronectin patterns. Four different cell categories are distinguished. (i) WT: Wildtype cells with vimentin. (ii) WT LatA+Noc: WT cells treated with latrunculin A and nocodazole to freshly polymerize microtubules while actin filaments are disrupted. (iii) VimKO: Vimentin knockout cells without vimentin. (iv) VimKO LatA+Noc: VimKO cells treated with latrunculin A and nocodazole. (a) Merged fluorescence images of microtubules (MT) in cyan, actin filaments in yellow, and vimentin IFs in magenta. (b) Fluorescence images of microtubules (MT). (c) Fluorescence images of actin filaments. In the drug treated cells WT LatA+Noc (ii) and VimKO LatA+Noc (iv), the normal actin architecture is disrupted. (d) Fluorescence images of vimentin IFs. As expected, in VimKO cells (iii) and VimKO LatA+Noc cells (iv) there is no vimentin present. Note that even within the same cell category, there are large cell-to-cell variations of network structures for each cytoskeletal filament type. Scalebars: 10 *µ*m.

### The alignment of microtubules is robust against the absence of vimentin and actin filaments

To obtain more insight into the typical microtubule network structure and its dependence on the surrounding actin and vimentin networks, we analyze the average alignment and orientation of microtubules in a spatially resolved manner with FFT on sliding windows.^29^ The alignment ranges between 0 for no alignment and 1 for perfectly parallel alignment and is represented in color-coded maps in Fig. 2a. Microtubules in all four cell categories show strong alignment in the cell interior as well as along the cell edge. These two regions are separated by a thin area of lower alignment. The average main orientation of microtubules in the interior follows a radial “wheel-like” structure from the center to the periphery and then transitions to a direction in parallel to the cell edge as displayed in color coded orientation maps in Fig. 2b. The low alignment area between cell interior and cell edge corresponds to the region where microtubules change from radial to azimuthal orientation. Both the alignment maps and orientation maps reveal an identical behavior of all cells with and without vimentin as well as with and without actin disruption. This analysis suggests that the global microtubule network structure is very robust and develops in a manner that is independent of vimentin IFs and actin filaments.

**Figure 2.**
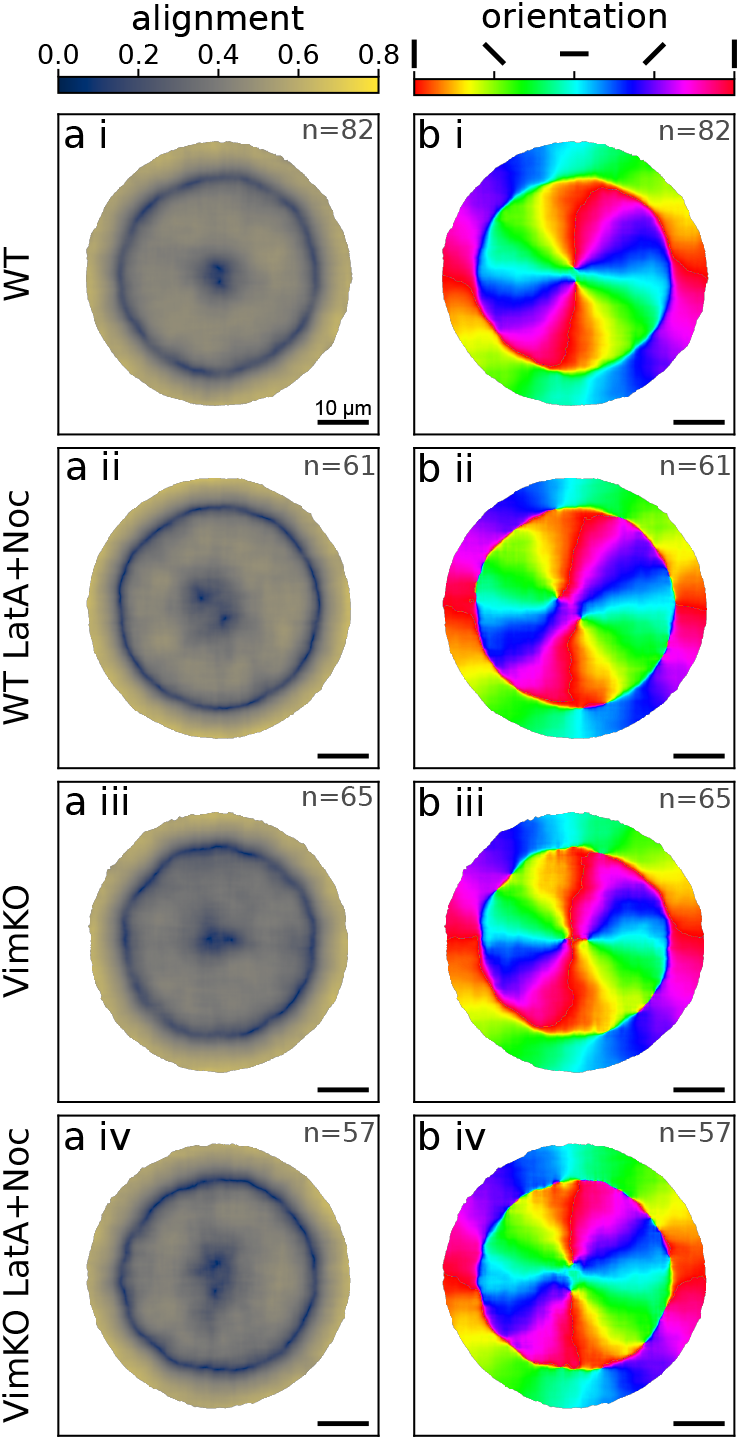
Influence of vimentin IFs and actin filaments on the structure of the microtubule network. (a) The averaged alignment of microtubules determined via Fourier analysis in circular cells. 0 corresponds to no alignment, 1 to complete alignment. (b) The averaged main orientation of microtubules with red and cyan indicating vertical and horizontal orientation, respectively. On the outer rim of the cells, the microtubules are oriented in parallel to the cell edge. In the interior, the microtubules are radially oriented. (i-iv) Four different cell categories with/without vimentin and with/without intact actin filaments. There are no differences in alignment and orientation between the different categories. Scalebars: 10 *µ*m. The number of cells over which we average is indicated in the top right of each panel.

### Actin and vimentin lead to small differences in average microtubule curvature

Despite displaying an overall reproducible average orientation in cells, microtubules appear highly curved. Bends and buckles have been attributed to non-thermal forces acting on microtubules in various manners^15,16^ and their mechanical response to such forces is believed to depend on the elastic properties of their local surrounding, including actin and vimentin networks.^17,18^ To investigate further how the surrounding cytoskeleton affects the bending of microtubules in cells, we analyze the curvature of automatically extracted microtubule segments. Coordinates of microtubules are extracted with the software SOAX.^30,31^ To minimize the influence of extraction artifacts in dense areas, we only take into consideration the center parts of segments between crossing points which are longer than 1.56 *µ*m as they are shown in Fig. 3a and b. The local curvature at each point of an extracted filament is determined by fitting a circle to its surrounding filament points as illustrated in Fig. 3c.

**Figure 3.**
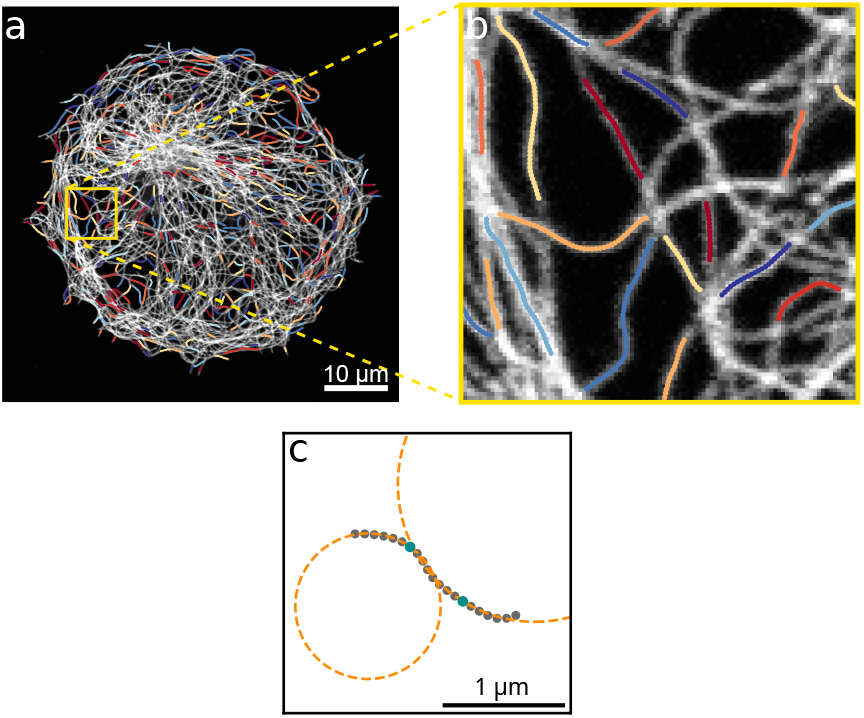
Curvature analysis of microtubules. (a) Microtubule segments which are used for curvature analysis overlaid on the fluorescence image of microtubules. The coordinates of the microtubule segments are extracted with SOAX and only the center parts of filaments with a length larger than 20 pixels are analyzed with respect to their curvature. (b) Enlargement of the yellow marked region in a. (c) Example filament for which the curvatures at the two example pixels marked in cyan are determined by fitting a circle to the local surrounding filament pixels.

The microtubule curvature values are averaged for each individual cell and we compare the distributions of cellular curvature averages in each cell category in Fig. 4a. For WT cells, the median cell average of microtubule curvature, rounded to the second decimal place, is 0.44 *µ*m^−1^ compared to 0.47 *µ*m^−1^ for VimKO cells. For the drug treated cells WT LatA+Noc and VimKO LatA+Noc, we obtain 0.46 *µ*m^−1^ and 0.50 *µ*m^−1^, respectively, as the median value of cellular average of microtubule curvature. Comparing WT (red) with VimKO (blue) and WT LatA+Noc (yellow) with VimKO LatA+Noc (green) yields a slightly higher cellular average of curvature for cells without vimentin (blue, green) in both cases. The comparison of WT with WT LatA+Noc and VimKO with VimKO LatA+Noc yields a slightly higher average cell curvature for the cells with actin disruption (yellow, green) in both cases. The differences of these four pairwise comparisons are on the order of only 4 to 7 % but are statistically significant at an overall significance level of 5% (Kruskal-Wallis test and Dunn’s test). We thus observe a small systematic trend of a slightly higher average microtubule curvature per cell for cells without vimentin and/or a disrupted actin network as compared to cells with vimentin and unperturbed actin.

**Figure 4.**
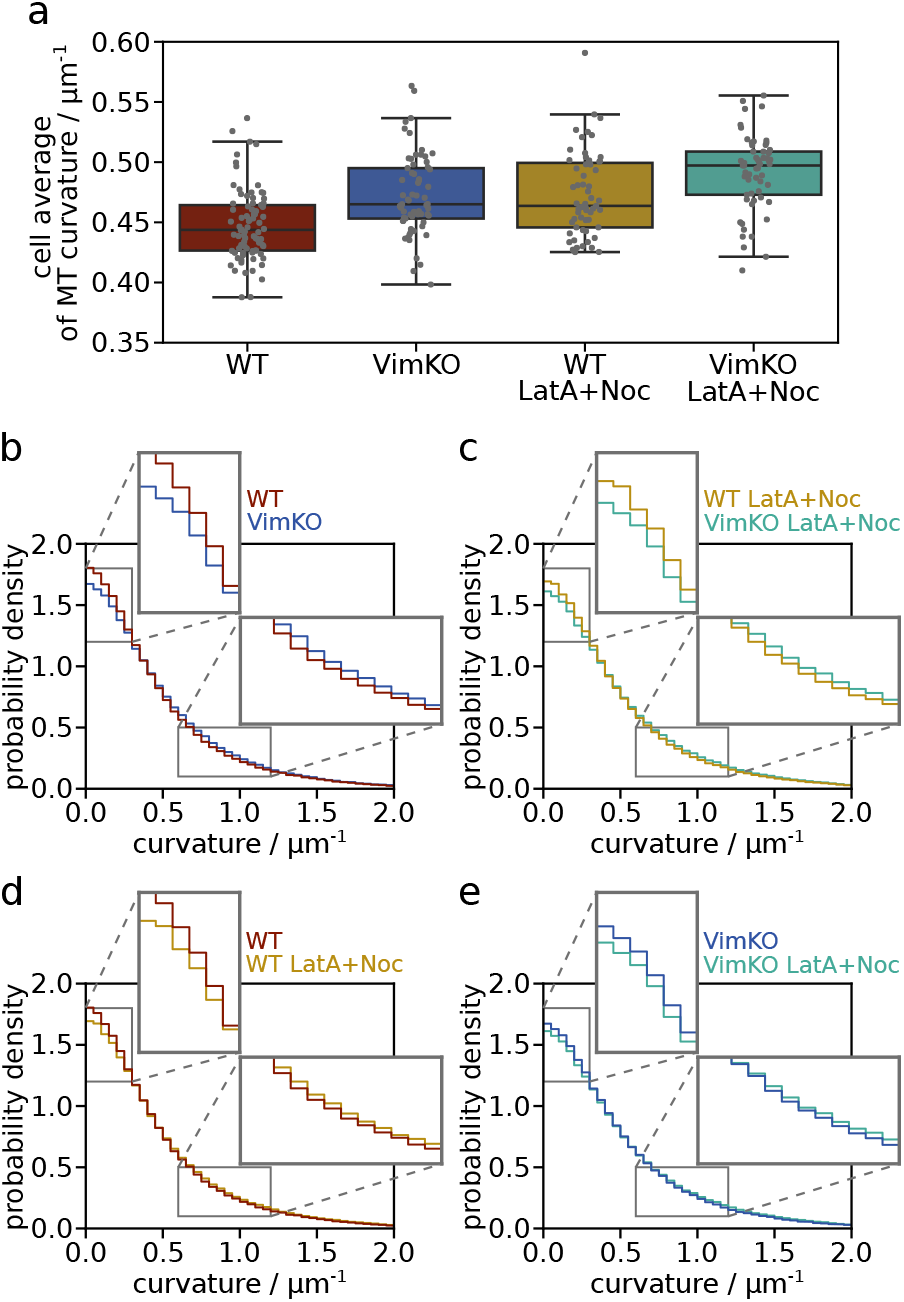
Influence of vimentin IFs and actin filaments on the average local microtubule curvature. (a) Distribution of cellular averages of the microtubule curvature for each cell category (with/without vimentin and with/without actin disruption). The average of all curvature values of one cell is represented by a gray dot for each cell. The boxplot of the distribution of all cellular averages shows the median and the interquartile range. (b, c, d, e) Histograms of the distribution of all curvature values in each category (irrespective of individual cells) with enlargements of a low-curvature range (0 to 0.3 *µ*m^−1^) and a higher-curvature range (0.6 to 1.2 *µ*m^−1^). Each histogram compares two categories: b) WT and VimKO cells, c) WT LatA+Noc and VimKO LatA+Noc cells, d) WT and WT LatA+Noc and e) VimKO and VimKO LatA+Noc. The bin width is 0.05 *µ*m^−1^. The comparison of cells with and without vimentin and the pharmacological disruption of actin filaments reveals a small shift to higher microtubule curvatures when cells are without vimentin and/or without intact actin filaments.

Since the differences are on such a small order and averaging per cell might obscure effects, we take a closer look at the entire distributions of curvature values without distinguishing between individual cells. Fig. 4b-e present histograms of the entire distributions of all curvature values within one cell category. Each histogram shows two distributions to compare between cells with and without vimentin (b and c) and cells with and without actin disruption (d and e). The distributions are all highly similar. We interpret this as a sign of reproducible mechanisms that make microtubules bend. The enlargements of the low and high curvature regimes make small differences visible: For each cell category comparison, one category has a higher proportion of low values and the other one shows a slightly higher proportion of values in the high curvature regime. The trend is such that in the comparison between WT and VimKO (both with and without drug treatment), the curvatures in VimKO cells are shifted towards higher values. Similarly, in the comparison between untreated and drug treated cells (for both WT and VimKO), the curvatures in cells with actin disruption show higher values. This behavior agrees with the comparison of cellular curvature averages shown in Fig. 4a and thus confirms a small trend of higher average curvatures in cells without vimentin and actin. It is not clear yet if these small differences are indeed mechanically induced by the presence and absence of vimentin and actin filaments. Therefore, we take a closer look at the curvature in a regional analysis.

### There is no direct link between average microtubule curvature and actin/vimentin density

We refine our curvature analysis and distinguish between regions with a higher and lower amount of vimentin or actin filaments. The heatmap of the averaged vimentin fluorescence intensity in WT cells in Fig. 5a shows the typical distribution of vimentin with a high density in the center that decreases towards the periphery of the cell. In contrast, the actin density is high in the periphery of the cells compared to only very few occasional fibers in the center, as visible in Fig. 5b. We divide each cell individually according to its shape into two regions, a center (cyan) and a rim (orange) region, as shown in an example in Fig. 5c. With this mask, we average the microtubule curvature in each region separately and compare it pairwise between cells with and without vimentin, or with and without actin disruption in Fig. 5d. The mean curvature is presented with a 95% confidence band as error bar which is in such a small range that it is barely visible. In Fig. 5d i and ii, we compare WT with VimKO and WT LatA+Noc with VimKO LatA+Noc cells, respectively. If the microtubule curvature were directly correlated with the density of vimentin, we would expect the difference between the cell categories to be larger in the center (cyan shaded) than in the rim region, since the density of vimentin is higher in the center thus creating a larger distinction between WT and VimKO cells. However, we observe a small and similar difference in both center and rim region for the comparison of WT and VimKO cells without drug treatment (Fig. 5d i) and a difference in the rim but not in the center for the comparison between WT LatA+Noc and VimKO LatA+Noc (Fig. 5d ii). These comparisons thus suggest that the microtubule curvature is not directly correlated with vimentin density. In the same manner, we compare cells with intact and disrupted actin filaments in Fig. 5d iii and iv. Here, if the microtubule curvature were directly dependent on the actin density, the difference between the two compared categories would be expected to be higher in the rim region (orange shaded) than in the center. This is indeed the case for VimKO and VimKO LatA+Noc (Fig. 5d iv) but not for WT and WT LatA+Noc (Fig. 5d iii) where there is a similar difference in both regions. Due to the missing consistency between WT and VimKO cells, it seems unlikely that there is a direct correlation between actin density and microtubule curvature. Therefore, we assume that the small differences in microtubule curvature that we observe between different cell categories in the different averaging approaches (comparison of cell average, entire distribution and regional average) stem from rather indirect effects and not from a direct mechanical impact of the actin and vimentin networks.

**Figure 5.**
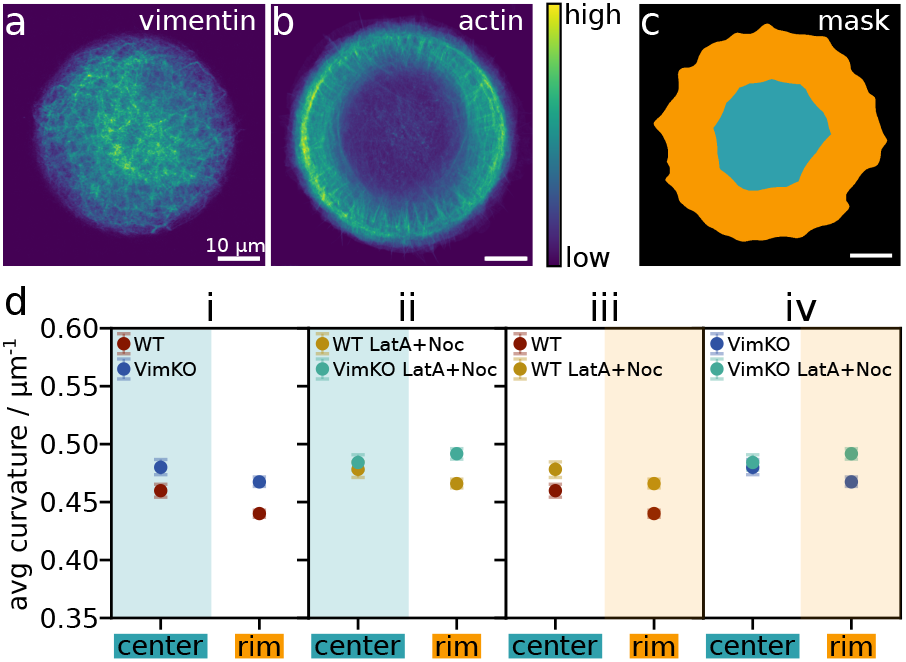
Regional analysis of microtubule curvature. (a) Heatmap of normalized vimentin IF fluorescence intensity averaged over 26 WT cells. (b) Heatmap of normalized actin fluorescence intensity averaged over 26 WT cells. (c) Example of a cell divided into two regions: center (cyan) and rim (orange). Each cell is individually divided according to its shape. Scalebars: 10 *µ*m. (d) Comparison of average microtubule curvature in the center and rim region for different cell categories: (i) WT and VimKO cells, (ii) WT LatA+Noc and VimKO LatA+Noc, (iii) WT and WT LatA+Noc, (iv) VimKO and VimKO LatA+Noc. For each pairwise comparison, the region with the colored background marks the region of the two where we would expect the greater difference between the categories if the microtubule curvature were directly influenced by the density of vimentin or actin. The mean curvature is presented with a 95% confidence band as error bar.

A repetition of the experiments and analysis with NIH3T3 mouse fibroblasts with vimentin and NIH3T3 cells, in which vimentin has been knocked out by CRISPR/Cas9 genome editing, confirms that there is no direct link between vimentin on the one hand and the global microtubule orientation and the local microtubule curvature on the other hand. The data and information on the NIH3T3 experiments are presented in the Supplementary information. Also in these cells, the microtubules are robustly oriented and aligned in a radial fashion independent of the presence and absence of vimentin. The local curvature of microtubules is very similar between all categories as well. For the drug treated NIH3T3 and NIH3T3 VimKO cells, the comparison yields the same trend as for the MEF cells. However, in contrast to the MEF VimKO cells, the NIH3T3 VimKO cells without drug treatment show a slightly lower instead of higher curvature than the corresponding WT cells. This inconsistency supports the conclusion that the small differences in curvature that we observe cannot be attributed to a direct effect of vimentin in the cell.

## Discussion

The interplay of microtubules, actin filaments and IFs in cells has found increasing interest but the picture of how cytoskeletal filaments influence each other remains complex with many details that are not clear yet. Here, we systematically study the influence of vimentin and actin networks on the global orientation and local curvature of microtubules in MEFs. The effect of vimentin in the cells is probed by comparing WT to VimKO MEFs and the effect of actin is probed by pharmacological disruption of the network. Micropatterning is employed as a powerful tool to compare cells with each other and to find average trends despite the natural large cell-to-cell variability.

We find that microtubules in circular cells are aligned and oriented in a radial manner in the interior, whereas in the periphery they are oriented in parallel to the cell edge. This alignment pattern is robust and independent of the presence or absence of vimentin and actin. The influence of actin was already investigated on triangular rat2 fibroblasts:^10^ Actin stress fibers guide microtubules to grow towards corners and focal adhesions, whereas they grow radially without actin fibers. Our result of the radial microtubule array being independent of actin filaments is thus in line with the previous study. It has been less clear, however, what the influence of vimentin is. Coalignment of microtubules and vimentin IFs has been found in different types of cells.^33–35^ Specifically in migrating RPE1 cells, vimentin filaments can serve as “tracks” for repolymerizing microtubules.^11^ Furthermore, in mouse myocytes, desmin IFs which form structural bundles with the Z-disc were found to mediate a more organized microtubule network as compared to a chaotic one in desmin knockout cells.^36^ The centrosome as the microtubule organizing center is assigned the function of controlling microtubule nucleation and is thus considered a main factor for the creation of a radial microtubule array.^37^ It has been previously shown that it is affected by the absence of vimentin in mouse fibroblasts: Its position displays a higher variability in polarized cells^32^ and it has a decreased size as well as a lower rate of microtubule-nucleation in VimKO MEFs compared to WT MEFs.^38^ Altogether, these findings hint to IFs, specifically vimentin IFs, impacting the structure of the microtubule network. Within this context, it is even more remarkable how robust and independent of vimentin the microtubule network orientation in the MEFs of our study is. We are cautious in generalizing this result to be valid across different cell types as the interactions between cytoskeletal filaments might depend strongly on the specific cell line.

On a much smaller length scale, we also analyze the local curvature of microtubules which yields information about the mechanical response of microtubules to forces. Our comparison of curvatures is based on large numbers of values from an automated analysis. The great spread of values demonstrates the need for large statistics and the need to average to detect overall trends within the large natural variability. Comparing the cellular average curvature as well as comparing the entire distribution of curvatures reveal a high similarity between all our investigated cell categories hinting to reproducible mechanisms that cause the bending of microtubules. Small differences on the order of few percent reveal that for MEFs, the microtubule curvature is on average slightly higher in cells without vimentin and without an intact actin network. Our regional analysis, however, indicates that the microtubule curvature does not directly correlate with vimentin or actin density. Therefore, we assume that the small curvature differences we observe are not a direct mechanical consequence of the vimentin and actin filament surrounding but are likely a rather indirect effect. This assumption is further supported by the result of the same analysis in NIH3T3 cells with vimentin and NIH3T3 cells with a CRISPR/Cas induced vimentin knockout. Comparing the microtubule curvatures in this case, the WT cells have a slightly higher curvature than the VimKO cells, thus displaying the opposite trend than the MEF cells even though both cells are mouse fibroblasts. Our study suggests that the average curvature of the large ensemble of microtubules in a mouse embryonic fibroblast is mostly independent of its surrounding vimentin and actin networks.

This conclusion may seem unintuitive considering previous studies but caution is necessary to distinguish between different observed phenomena. It has been widely accepted that the buckling of microtubules under longitudinal compression is affected by their viscoelastic surrounding.^17,18,39^ With a higher elastic modulus of the surrounding, the buckling wavelength decreases according to the constrained buckling theory. Thus, since both actin and vimentin networks generally make the cytoplasm stiffer,^19–21,40^ a larger microtubule buckling wavelength is expected when vimentin and actin are lacking in a cell.^18^ This was demonstrated with microtubules in Cos7 cells which were compressed with a microneedle and displayed indeed an increased buckling wavelength when actin filaments were depolymerized with cytochalasin D.^18^ Also in HeLa cells, localized bends of microtubules showed an increase upon actin filament disruption.^41^ An important difference distinguishes these studies from ours: Whereas the other studies investigated individual, manually selected microtubules with dynamic buckling, we investigate a large ensemble of automatically extracted filaments in static images of whole cells. So we have, on the one hand, single, strong buckling events and, on the other hand, the general bending of microtubules that occurs everywhere and to different degrees. For the analysis of microtubule bending in static images, the picture remains complex. A previous study of cryo-electron tomograms of HeLa and P19 cells came to the conclusion that microtubules are more curved when they are entangled with actin filaments and IFs which is in line with the constrained buckling theory, yet the throughput of this study was only a few cells.^23^ Interestingly, when repolymerizing microtubules while actin filaments are disrupted with latrunculin B, the persistence length of microtubules in the tomograms was slightly lower than normally, which is in contrast to the other results. This finding is comparable to our finding in drug-treated cells when the microtubule curvatures are slightly higher after microtubules have repolymerized during actin disruption. In contrast, in living Xenopus laevis melanophores, the persistence length of microtubules was not observed to be affected by the disruption of actin with latrunculin B.^24^ However, upon the depletion of vimentin, the persistence length was decreased which corresponds to higher curvatures and contrasts the studies of simple disruption of actin filaments.^24^ These differences underline the complexity and diversity of the relationship between microtubule curvature and the cytoskeletal surrounding which might strongly depend on the cell type. For the cell type we use here, i.e., MEFs from WT and VimKO littermate mice, another study described abnormal microtubule shapes and larger wavelength buckling in VimKO cells, yet found a substantially larger average curvature in VimKO cells.^42^

Altogether, different studies with different cells and methods yield different results on the influence of vimentin and actin networks on microtubule shapes. An important distinction has to be made between the analysis of particularly dynamically buckling microtubules and the analysis of the static shape of microtubules without specific selection. The former case of dynamically buckling microtubules might demonstrate well the constrained buckling theory in which a compressed microtubule is supported by the surrounding cytoskeletal networks, but the large ensemble of microtubules might experience a variety of different scenarios leading to their overall curvature. Nevertheless, also for the static, bent shape of microtubules, an important role has been attributed to the elastic surrounding: A previous study found that the large-scale bent trajectory of a microtubule is primarily created while the microtubule grows and is stabilized in time afterwards, presumably by the elastic surrounding and in particular by the cytoskeleton.^18^ Considering this point of view, the independence of the microtubule curvature from vimentin and actin networks in our cells means that the majority of bending is maintained by cellular structures other than the cytoskeleton. Microtubule motors, crosslinkers or crossing points with other microtubules could be possible candidates to serve as anchoring points for the microtubules and thus yield a similar effect as a continuous elastic support. For example, an in vitro gliding assay with microtubules and motors but no other cytoskeletal filaments exhibited curvatures in a similar range to those in cells.^16^ Also, the compression of microtubules linked with kinesin motors to a PDMS substrate showed that the buckling wavelength decreased with an increased kinesin concentration. ^43^ These studies demonstrate that bent trajectories of microtubules do not necessarily require a cytoskeletal surrounding. The cytoskeleton might function as a continuous elastic matrix on the larger scale of the whole cell. However, depending on its network properties such as mesh size and density which can vary between cell types and states, it could possibly be too inhomogeneous on a local scale to serve as a continuous elastic surrounding of microtubules. Our study suggests that the role of vimentin and actin on microtubule bending in unpolarized mouse embryonic fibroblasts might be unexpectedly small and of rather indirect nature. The complex interactions between the cytoskeletal filaments could be cell type dependent for instance due to different network architectures and motor activities, and may vary with respect to an unpolarized or migrating state of the cell. Overall, in unpolarized MEFs, the microtubule system appears robust in both its global alignment and orientation as well as its local mechanical response to forces. It might be more independent of other cytoskeletal filaments than expected from the interactions found in other cell types.

## Acknowledgement

We thank Charlotta Lorenz, Anna V. Schepers, Jan Christoph Thiele, Alexandre Schaeffer and Gilles Mordant for fruitful discussions. We thank John Eriksson for providing the MEF WT and MEF VimKO cells. This work was funded by the Deutsche Forschungsgemeinschaft (DFG, German Research Foundation): Project-ID 432680300 - SFB 1456/A04 (to S.K. and A.M.); Project-ID 449750155 - RTG 2756/A2, A7 (to S.K.); Project-ID-390729940 - in the framework of Germany’s Excellence Strategy, EXC 2067/1 (to S.K. and A.M.). The work was further financially supported by the European Research Council (ERC, Grant No. CoG 724932, to S.K.).

## Author contributions

Conceptualization: L.S. and S.K.; formal analysis: A.B. and D.V.; funding acquisition: A.M. and S.K.; investigation: A.B. and U.R.; software: A.B., D.V. and G.N.; supervision: A.M., L.S. and S.K.; visualization: A.B. and S.K.; writing - original draft: A.B. and S.K.; writing - review & editing: all authors

