## Supplementary Information for "Global alignment and local curvature of microtubules in mouse fibroblasts are robust against perturbations of vimentin and actin"

### Additional details of data analysis

#### Image preprocessing

Every image is pre-processed following the workflow shown in Fig. S1. To ensure the same position of each cell, every image is spatially shifted such that the center of mass is located in the center of the image. This shift is determined with the aid of a segmentation of the interior of the cell. To this end, each microtubule image is uniquely background-corrected by subtracting the minimum intensity value. A Gaussian filter is applied and the image is binarized by thresholding with Otsu's method. Any small holes remaining in the segmented cell area are filled and a binary erosion is applied. The resulting binary image is shifted to position the center of mass at the same pixel for all images. This aligned, segmented cell image $I_{\mathrm{segm}}$ is used as an auxiliary image for further analysis steps. The same shift that is used for the segmented image is applied to the fluorescence image to obtain an aligned fluorescence image as the starting point for all further analysis.

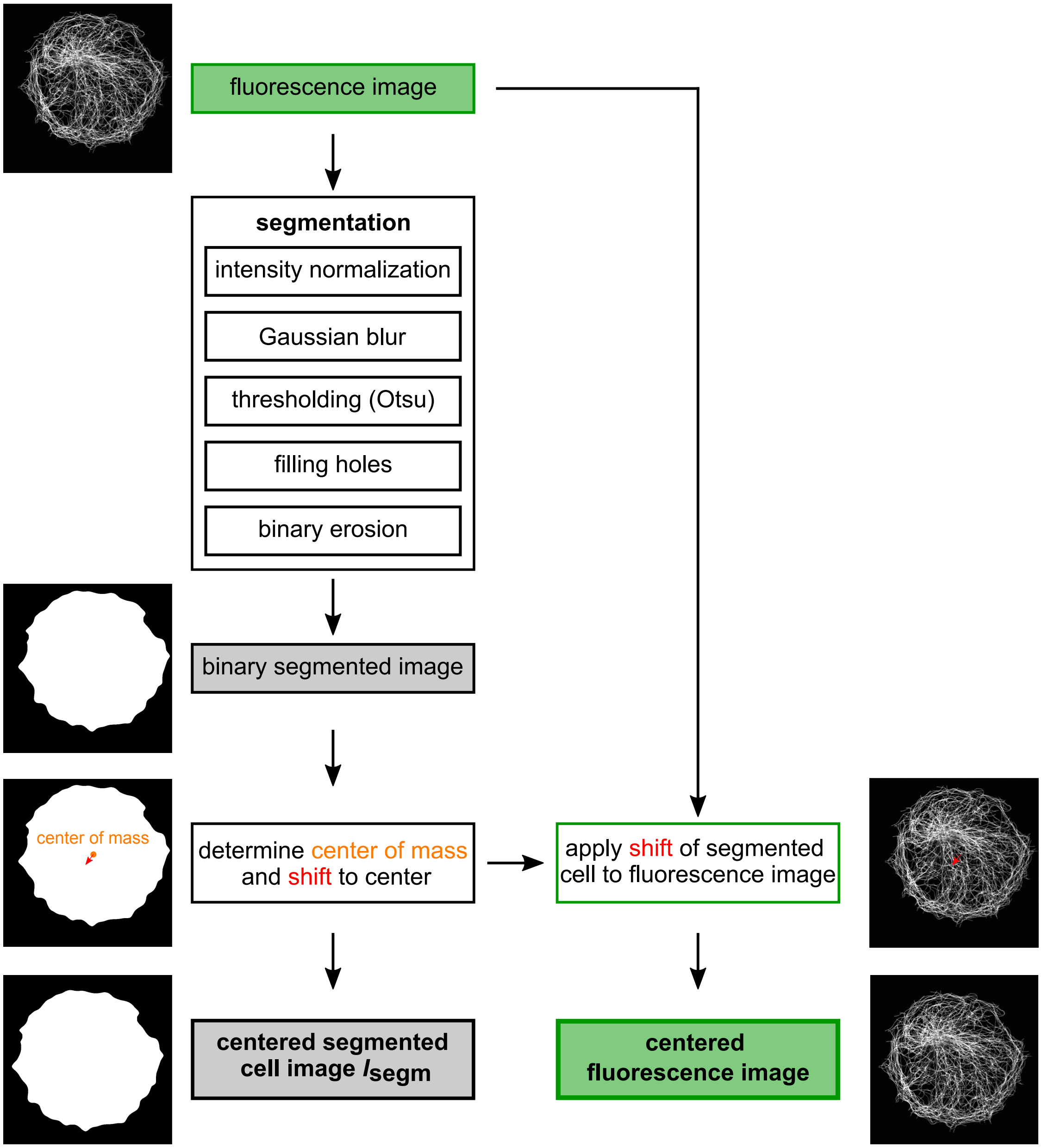


Fig. S1: Workflow for preprocessing of all fluorescence images. All cells are centered in the image and the aligned binary image $I_{\mathrm{segm}}$ is used as an auxiliary image for further analysis steps.

#### Binary cell area mask

A binary mask for the typical area of cells is created by overlaying all segmented cell images $I_{\text{segm}}$ as schematically illustrated in Fig. S2. To exclude outliers of unusually large cell shapes, only pixels belonging to more than 10% of all the segmented cells constitute together the binary cell mask. This mask is applied in the end to the averaged orientation and alignment map of all cells.

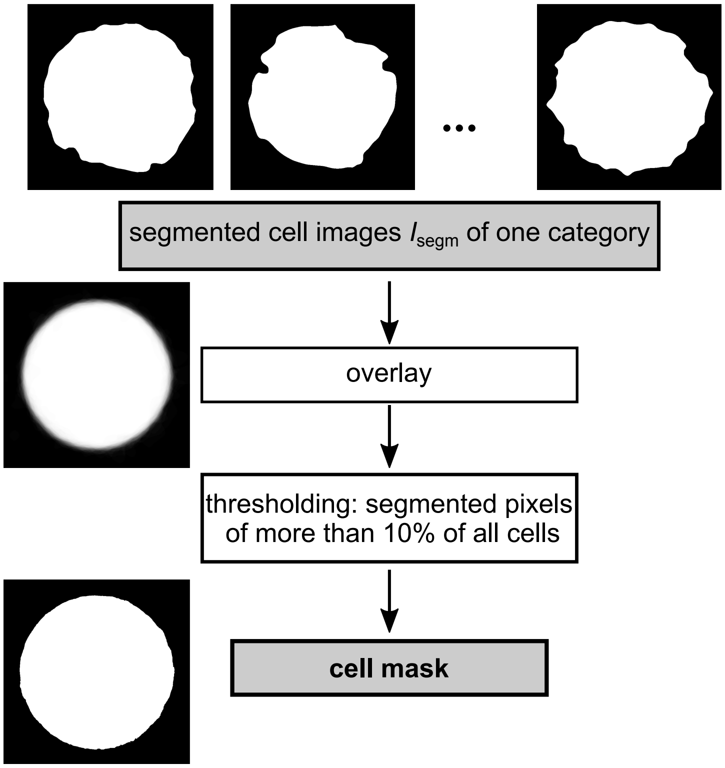


Fig. S2: Workflow of processing steps to create a binary mask for cells of one category.

#### Orientation analysis via Fourier transform

The average local alignment and orientation of microtubules in the circular cells is analyzed with Python code based on the AFT tool (Alignment by Fourier Transform).[1] The schematic workflow for the analysis is depicted in Fig. S3. In brief, an FFT is calculated for a sliding window of size 71x71 pixels for each cell. For each window position, the FFT amplitude images are then averaged over all cells and a principal component analysis (PCA) is conducted to obtain eigenvectors and eigenvalues. The direction of the eigenvector with the largest eigenvalue determines the main orientation of the microtubules and the values of the largest eigenvalue $\lambda_1$ and the smallest eigenvalue $\lambda_2$ are used to calculate the eccentricity of the FFT signal as $e = \sqrt{1 - \lambda_2/\lambda_1}$. This eccentricity serves as a measure for the degree of microtubule alignment with 0 indicating no alignment and 1 indicating maximum alignment.

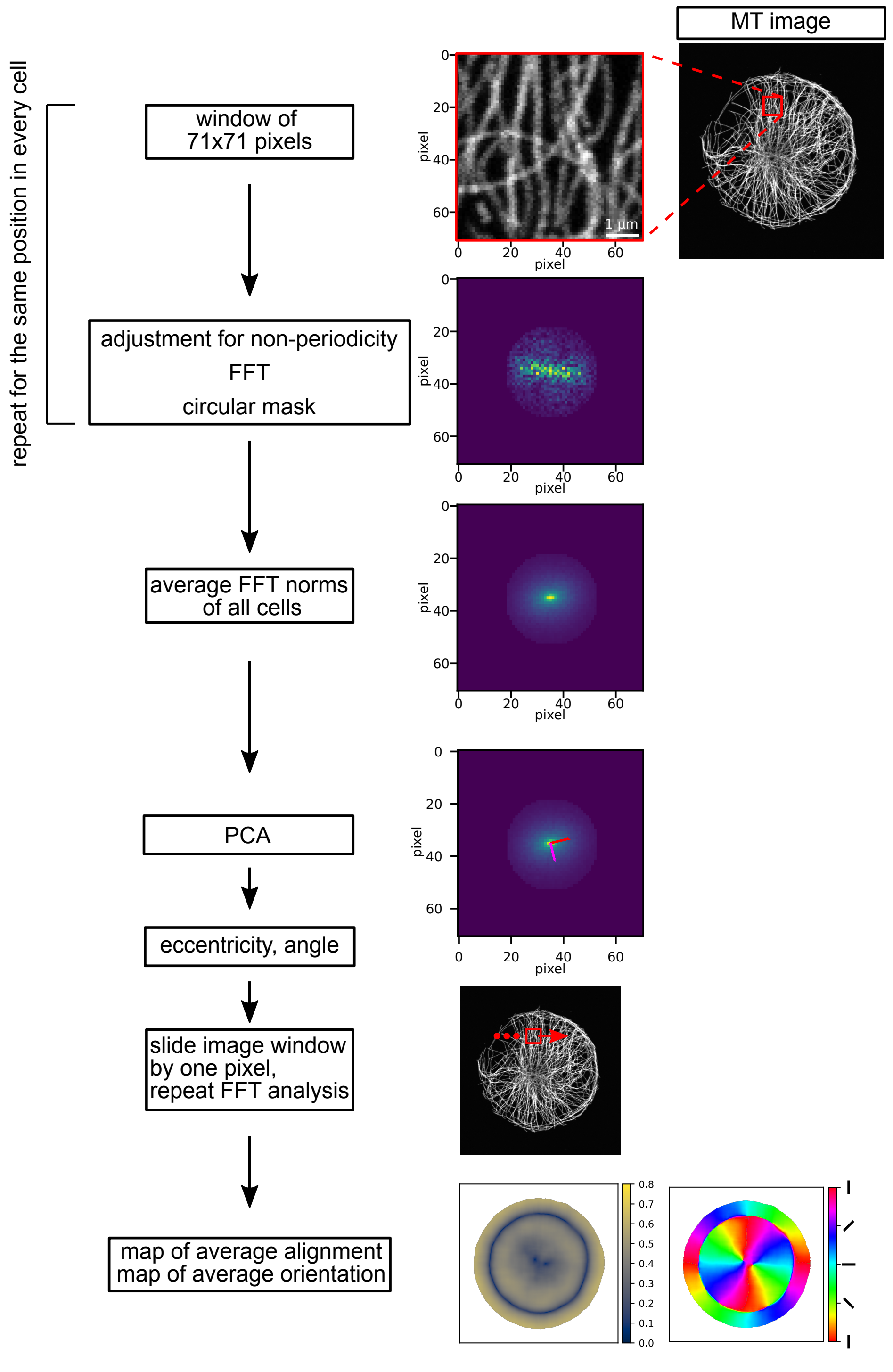


Fig. S3: Schematic workflow for orientation analysis. A sliding window of 71x71 pixels is applied to the image of microtubules (MT image). The subwindow undergoes a correction for non-periodicity to avoid artifacts in the following fast Fourier transform (FFT). A circular mask is applied to the norm of the FFT to filter out high frequency noise. This FFT procedure is applied to the same window of every cell image. The FFT norms for one window position are averaged and a PCA is conducted. The red arrow in the corresponding image marks the direction of the eigenvector with the largest eigenvalue, the pink arrow the eigenvector with the smallest eigenvalue. With the PCA, an eccentricity value as a measure for the alignment and an angle as a main orientation is obtained. This workflow is repeated for every window position sliding along the original image in steps of 1 pixel. The result is a map of average alignment and a map of average orientation.

#### Calculation of confidence band as error bar in regional curvature analysis

We quantify the variability of the estimated regional curvature by providing pointwise confidence intervals for the population counterpart of the mean curvature $\bar{c}$, which can be thought of as the limit value we would obtain if we had access to data from an infinite number of cells. We can determine each confidence interval using a cluster bootstrap procedure.[2] This procedure consists in sampling $g$ times with replacement from the detected filaments (clusters) $F_1, \ldots, F_g$ (with $g \in \mathbb{N}$ being the total number of the detected filaments) instead of sampling from the filament points. This clustered resampling method has the advantage of taking into account the strong correlation between curvature values of points that belong to the same filament. After constructing $B = 1\,000$ resampled datasets, we compute the corresponding bootstrap replicas $\bar{c}^{(j)}$, with $j = 1, \ldots, B$. We can then determine a basic bootstrap confidence interval $I \subset \mathbb{R}$, also known as the reverse percentile interval, given by:

$$I = (2\bar{c} - c^*_{(1-\alpha/2)},\, 2\bar{c} - c^*_{(\alpha/2)})$$

where $c^*_{(1-\alpha/2)}$ denotes the $1 - \alpha/2$ percentile of the bootstrapped replicas $\bar{c}^{(1)}, \ldots, \bar{c}^{(1000)}$. We set $\alpha = 0.05$ to calculate 95% pointwise confidence intervals.

### Analysis of NIH3T3 cells

In addition to MEF WT and MEF VimKO cells from two littermate mice, we also analyze NIH3T3 cells with vimentin and NIH3T3 cells in which vimentin is knocked out via CRISPR/Cas9 genome editing. We refer to these VimKO cells as NIH3T3VimKO. The experimental procedure is the same as for the MEF WT and MEF VimKO cells with minor differences as described below.

#### Micropatterning for NIH3T3 cells

For NIH3T3 cells with vimentin and without drug treatment, the micropatterning of the glass substrates with the photopatterning system PRIMO is performed with the photoinitiator PLPP liquid instead of PLPP gel according to instructions of Alvéole. In brief, PDMS stencils are placed on the plasma-treated glass surface and the well is filled with 0.1 mg/ml PLL-PEG (PLL(20)-g[3.5]-PEG(2), SuSoS AG, Dübendorf, Switzerland) in PBS as passivation reagent and incubated for 1h. The wells are rinsed with PBS followed by the addition of PLPP liquid. The UV dose for illumination with PRIMO when using PLPP liquid is 1500 $mJ/mm^2$. After the illumination, the sample is washed with PBS and fibronectin at a final concentration of 50$\mu$g/ml in PBS is incubated for 1 to 2h. Unbound fibronectin is washed off with PBS. For NIH3T3VimKO cells and NIH3T3 cells with drug treatment, the micropatterning is performed with PLPP gel according to the description in Materials and methods of the main text.

#### Cell culture and experimental procedure

NIH3T3 cells are purchased through DSMZ (ACC-59-NIH-3T3 fibroblasts, Leibniz Institute DSMZ - German collection of Microorganisms and Cell Cultures GmbH, Braunschweig, Germany). From these NIH3T3 cells, vimentin knockout cells (NIH3T3 VimKO) are generated by CRISPR/Cas9 genome editing using Trueguide synthetic guide RNA (Invitrogen,

#A35533, ID#CRISPR492856_SGM), TrueCut Cas9 Protein v2 (Invitrogen, #A36496) and lipid-based transfection with Lipofectamine CRISPRMAX Cas9 Transfection Reagent (Invitrogen, #CMAX00008) according to the manufacturer's instructions. The result of this CRISPR/Cas9 transfection is a mixed culture of NIH3T3 cells with varying degree of vimentin knockout with which experiments are conducted and only cells with no vimentin, as identified by immunostaining, are selected for further analysis.

We perform the same experiment as for MEF WT and MEF VimKO to repolymerize microtubules while actin filaments are disrupted and refer to these cells as NIH3T3 LatA+Noc and NIH3T3VimKO LatA+Noc. Latrunculin A (Sigma-Aldrich, #428021) is used at a final concentration of 0.6 $\mu$M and nocodazole at a final concentration of 2 $\mu$M.

The fixation is identical to the fixation of MEF WT and MEF VimKO cells. Microtubules and vimentin IFs are immunostained. In total, we analyze 102, 77, 26 and 26 cells of the cell categories NIH3T3, NIH3T3VimKO, NIH3T3 LatA+Noc and NIH3T3VimKO LatA+Noc, respectively. We analyze the microtubule alignment and microtubule curvature in these cells in the same way as described for the MEF cells.

#### Alignment of microtubules in NIH3T3 cells

The alignment and orientation maps of microtubules in NIH3T3 cells with/without vimentin and with/without actin disruption are displayed in Fig. S4. They show the same structure as for the corresponding cell categories of MEF cells: The microtubules are aligned in a radial fashion from the center to the periphery and then change from their radial to an azimuthal orientation in parallel to the cell edge. For NIH3T3 LatA+Noc, the orientation "wheel" is less clear and more noisy than for the other categories but since the cell number of 26 is rather low, this effect could stem from natural cell-to-cell variation.

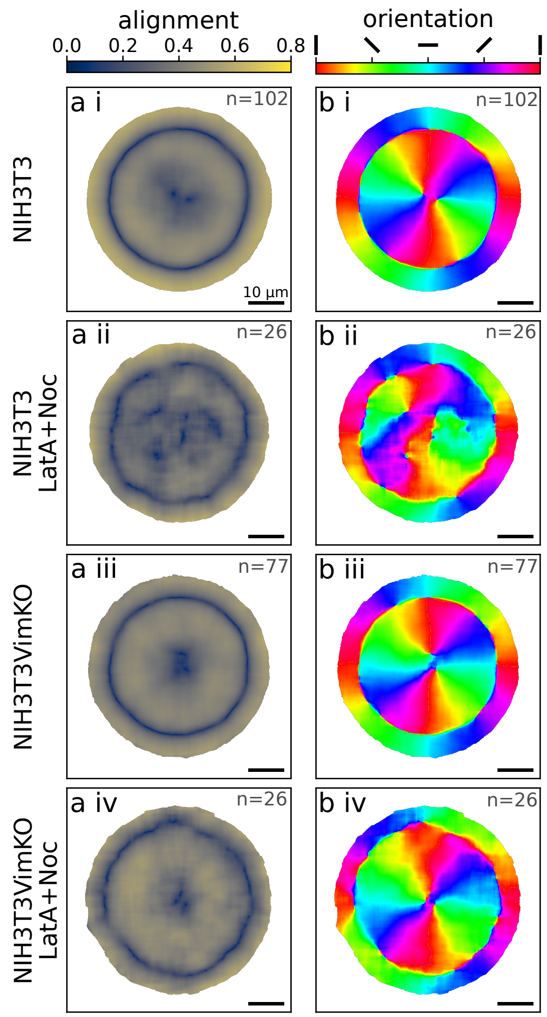


Fig. S4: Influence of vimentin IFs and actin filaments on the structure of the microtubule network in NIH3T3 cells. (a) The averaged alignment of microtubules determined via Fourier analysis in circular cells. 0 corresponds to no alignment, 1 to complete alignment. (b) The averaged main orientation of microtubules with red and cyan indicating vertical and horizontal orientation, respectively. On the outer rim of the cells, the microtubules are oriented in parallel to the cell edge. In the interior, the microtubules are radially oriented. (i-iv) Four different cell categories with/without vimentin and with/without intact actin filaments. a ii and b ii show more noisy maps, otherwise there are no fundamental differences in alignment and orientation between the different cells. The numbers of cells over which is averaged in each category is given in the top right of the plots. Scalebars: 10 $\mu$m.

#### Curvature of microtubules in NIH3T3 cells

The curvature analysis of microtubules in the different categories of NIH3T3 cells is presented with the cellular average and the entire distribution of curvature values in Fig. S5. As for MEF cells, we perform first an analysis of the cellular average curvature of microtubules by averaging all curvature values per cell and representing the distributions of cellular averages in boxplots in Fig. S5a. The NIH3T3VimKO cells have a slightly lower median of $0.44\,\mu\text{m}^{-}1$ (rounded to the second decimal place) compared to the NIH3T3 cells with $0.45\,\mu\text{m}^{-}1$ which is thus the opposite trend of MEF cells where the VimKO cells had a higher curvature median than the WT cells. However, comparing the drug treated NIH3T3 LatA+Noc and NIH3T3VimKO LatA+Noc cells yields a slightly higher curvature of $0.46\,\mu\text{m}^{-}1$ for the cells without vimentin than for the cells with vimentin ($0.44\,\mu\text{m}^{-}1$). Comparing the cells with and without actin disruption yields contrasting trends as well: The actin disruption slightly lowers the curvature median for the NIH3T3 cells whereas for the NIH3T3VimKO cells, the actin disruption increases the curvature median. These small differences between the four categories of NIH3T3 cells are not statistically significant, in contrast to the results of the comparison between MEF cells with/without vimentin and actin filaments. The comparison of the entire distributions of all curvature values without distinguishing between individual cells is presented in the form of histograms in Fig. S5b-e. The pairwise comparisons between cells with/without vimentin and with/without intact actin filaments fit with the small trends of the cellular curvature average in Fig. S5a.

The regional analysis of the microtubule curvature for which the cell is divided into a center and a rim region as shown in an example in Fig. S6b does not show a clear correlation between microtubule curvature and region. The vimentin density is much higher in the center region as visible in the heatmap of Fig. S6a, revealing a higher gradient in density as in the MEF WT cells. Yet, the difference between NIH3T3 with and without vimentin is not larger in the center region than in the rim region as visible in Fig. S6c. When comparing drug-treated cells with untreated cells, the difference varies between the center and the rim region but with opposite effects for NIH3T3 and for NIH3T3VimKO cells. Thus, there is no consistent difference that can be particularly attributed to a higher actin density in the rim region.

Overall, the comparison of the different categories of NIH3T3 cells does not yield a picture of a consistent connection between the microtubule curvature and vimentin IFs and actin filaments.

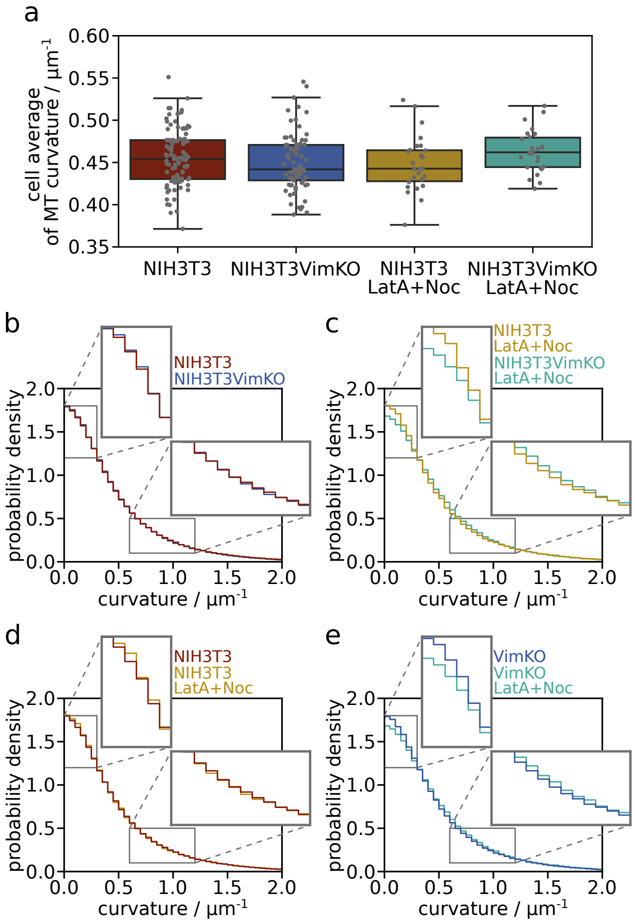


Fig. S5: Curvature analysis of microtubules in NIH3T3 cells. (a) Distribution of cellular averages of the microtubule curvature for each cell category (with/without vimentin and with/without actin disruption). The average of all curvature values of one cell is represented by a gray dot. The boxplot of the distribution of all cellular averages shows the median and the interquartile range. (b, c, d, e) Histograms of the distribution of all curvature values in each category (irrespective of individual cells) with enlargements of a low-curvature range (0 to $0.3\mu m^{-1}$) and a higher-curvature range (0.6 to $1.2\mu m^{-1}$). Each histogram compares two categories: NIH3T3 and NIH3T3VimKO cells in b, NIH3T3 LatA+Noc and NIH3T3VimKO LatA+Noc cells in c, NIH3T3 and NIH3T3 LatA+Noc in d and NIH3T3VimKO and NIH3T3VimKO LatA+Noc in e. In b and d, there is nearly no difference between the distributions, in c and e, the cells with actin filament disruption have slightly higher microtubule curvatures than the cells with intact actin filaments.

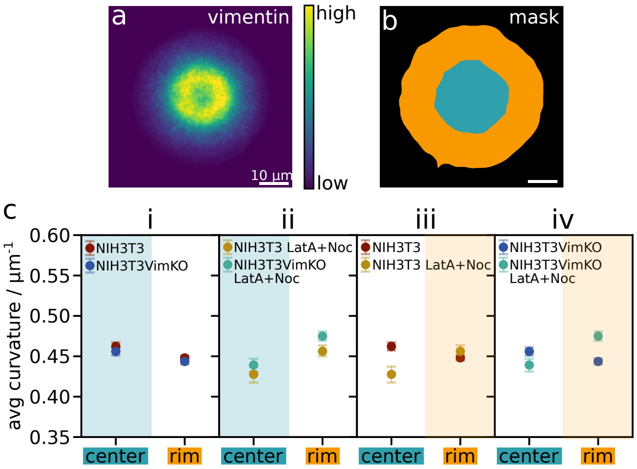


Fig. S6: Regional analysis of microtubule curvature in NIH3T3 cells. (a) Heatmap of normalized vimentin IF fluorescence intensity averaged over 102 NIH3T3 cells. (b) Example of a cell divided into two regions: center (cyan) and rim (orange). Each cell is individually divided according to its shape. Scalebars: 10 $\mu$m. (d) Comparison of average microtubule curvature in the center and rim region for different cell categories: (i) NIH3T3 and NIH3T3VimKO cells, (ii) NIH3T3 LatA+Noc and NIH3T3VimKO LatA+Noc, (iii) NIH3T3 and NIH3T3 LatA+Noc, (iv) NIH3T3VimKO and NIH3T3VimKO LatA+Noc. For each pairwise comparison, the region with the colored background marks the region of the two where we would expect the greater difference between the categories if the microtubule curvature were directly influenced by the density of vimentin or actin. The mean curvature is presented with a 95% confidence band as error bar.
